# Bone response to intermittent parathyroid hormone (PTH) is both genetic and sex specific in mice

**DOI:** 10.64898/2026.06.09.729629

**Authors:** Douglas J. Adams, Dana A. Godfrey, Shane E. Ridoux, Robert D. Maynard, Nicole S. Szeto, Ethan M. Lange, Cheryl L. Ackert-Bicknell

**Author notes:** Corresponding Author: Cheryl L Ackert-Bicknell 12800 E. 19th Avenue | Mail Stop 8343 Department of Orthopedics University of Colorado Aurora, CO, 80045.

## Abstract

Teriparatide (PTH 1-34) is an anabolic agent used to treat osteoporosis, yet clinical response varies widely among patients. To investigate genetic and sex-specific determinants of skeletal response, we administered intermittent PTH to male and female mice from eight genetically diverse inbred strains. Mice were treated for four weeks, and bone phenotypes were assessed via DXA, microCT, and mechanical testing. Response to PTH was highly strain- and sex-dependent, with some strains responding at the femur but not the spine, and vice versa. Heritability estimates for PTH-induced changes in areal bone mineral density (aBMD), cortical bone area at the femoral mid-diaphysis, femur strength, and trabecular bone volume fraction (BV/TV) ranged from moderate to high, with BV/TV showing the strongest genetic influence. Cortical bone response mechanisms differed by sex: males exhibited periosteal expansion, while females showed endosteal remodeling. These findings mirror clinical observations where non-response to PTH can be anatomic site specific. Further, our study suggests that genetic background and sex significantly influence bone response to PTH. Our data support the future use of genetically diverse mouse models to elucidate the genetic architecture of skeletal response to PTH.

**Highlights:**

- PTH response varied by genetics, sex, and skeletal site (femur vs spine).
- Trabecular bone volume showed the strongest genetic effect (heritability) with PTH treatment.
- Response to PTH was sex-specific.

## INTRODUCTION

Teriparatide is an anabolic agent used for the treatment of both primary osteoporosis in men and women(^1^) and secondary osteoporosis due to glucocorticoid use (^2,3^). It is a recombinant peptide consisting of the first 34 amino acids of parathyroid hormone (PTH)(^4^) and like native PTH, teriparatide can stimulate bone formation by osteoblasts and resorption by osteoclasts(^2^). Teriparatide is commonly administered once daily such that patients are exposed in an intermittent fashion, leading to an increase in osteoblast number and increased bone mass(^2,5^). Clinical trials data shows excellent efficacy for the reduction of vertebral fracture and non-vertebral fracture, as well as being an effective agent to increase bone mineral density (BMD) of the spine, hip and whole body(^2^). Other anabolic agents currently approved in the US include abaloparatide, which acts via a similar mechanism as teriparatide, and romozosomab, a monoclonal antibody that acts through the WNT signaling pathway (^4^).

Mounting evidence has suggested that not all patients respond to teriparatide (^6–8^) and while factors such as age and vitamin D status have been shown to contribute to the magnitude of response, a full understanding of why some patients do not respond is lacking(^8,9^). Modifiable environmental factors impact bone in an interactive manner with genetic factors. Indeed, many such Gene by Environment (G*E) interactions impacting bone have been described for factors such as dietary fat, exercise and vitamin D use(^10^). Like these common environmental variables, response to teriparatide is also thought to be partially mediated by genetic factors. A small genome-wide association study (GWAS) was conducted in humans (437 participants) for the phenotypes of “*response of BMD at either the lumbar spine or hip to teriparatide*” (^11^) and allelic differences at the *CXCR4* gene on human Chromosome (Chr) 2 were found to reach statistical significance for both BMD sites(^11^). However, a large and well powered Genome Wide * Environment Interaction Study (GWIS) for any drug in humans is very difficult and exceedingly costly. Further, the sample size needed to have statistical significance and avoid the pitfall known as a Winner’s Curse often exceeds that which is needed for a straightforward GWAS (^12,13^). This makes it very difficult to resolve the mechanisms by which genetic factors impact the effect of teriparatide/PTH on bone in humans. Such studies are substantially more feasible and cost effective in mouse models. To design such a study, the heritability of response to teriparatide/PTH needs to be known to determine study power. The relative heritability of bone compartment specific phenotypes needs to be known to efficaciously allocate resources in a GWIS. As many phenotypes cannot be collected in humans, and the only existing GWAS related to the topic of PTH response is small, heritability estimates are needed to move the field forward.

To fill this gap in knowledge, we treated male and female mice from eight inbred strains with either vehicle or PTH for four weeks to better define the heritability of teriparatide response. These 8 strains were chosen as they are the founder strains for two genetic mapping populations, the Diversity Outbred population (^14^) and the Collaborative Cross (^15^) recombinant inbred set. By using these 8 strains, we would be able to conduct the power calculations needed for a mouse GWIS study using the Diversity Outbred population in the future. We determined that the anatomic site of response to treatment varied widely among the strains and that there were strong sex differences among the strains. Further, we observed that some strains would only respond in the cortical compartment and others would present with trabecular response phenotypes. Our results reflect that which was seen in the human teriparatide response GWAS but provide valuable insight into the nature of response as we can include phenotypes that cannot be captured in humans.

## MATERIALS AND METHODS

### Animals and PTH treatment

All studies involving mice were reviewed and approved by the Institutional Animal Care and Use Committee (IACUC) of the University of Colorado (CU), Anschutz Medical Campus (Protocol number: 00902). All mice were fed Teklad Irradiated Global Soy Protein-Free Extruded Rodent Diet (Teklad, Cat no. 2920X) and had *ad lib*. access to food and water. The mice were maintained on a 14hr:10hr light:dark cycle. Mice used in this study were purchased from The Jackson Laboratory and the following 8 strains were used with strain stock numbers provided in parentheses: A/J (#000646 ), C57BL/6J (B6, #000664 ), 129S1/SvImJ (129, #002448), NOD/ShiLtJ (NOD, #001976), NZO/HlLtJ (NZO, #002105), CAST/EiJ (CAST, #000928), PWK/PhJ (PWK, #003715 ), and WSB/EiJ (WSB, #001145 ). For each strain, ∼40 mice were purchased, half of which were male. Mice were purchased at or near weaning age and allowed to acclimate to the CU vivarium until the age of 12 weeks. Each strain of mice was divided into two equal groups for daily i.p. injection, 5 days per week for 4 weeks, with either vehicle (saline) or PTH:1-34 (40 mg/kg; Bachem Product Number 4011474.0005). Group assignment was determined at the time of receipt from the vendor. Mice were randomly assigned to each group since each mouse of each strain is a genetic clone of all other mice of that strain(^16^). Differences in any phenotype within a strain and sex group can only arise from environmental factors and measurement error. All of these strains have been inbred by brother-sister matings for over 4 decades and all strains have been sequenced to 200X coverage to test for uniform homozygosity(^17^). All mice within a strain experienced the same environmental conditions, as we ordered all mice of a strain at once. While a higher dose for PTH is often used for in rodent studies, the concern was that a ceiling effect for trabecular bone increase could occur in strains with high bone volume fraction (BV/TV) with higher doses, creating an error in data interpretation. We chose this lower dose that has been used in other previous studies (^18–20^). At 12 weeks of age a baseline DXA scan was conducted. At 16 weeks of age all mice were euthanized and whole-body BMD was measured by DXA. The lumbar spine and left femur were collected, wrapped in saline-soaked gauze, and stored frozen at −20°C for microCT imaging (both) and mechanical testing (femur). A small number of samples were damaged during dissection and were not used for phenotyping. The number of mice per group for each type of measurement is shown in Supplemental Tables 1-6.

### Whole Body BMD and Body Composition via Dual X-ray Absorptiometry (DXA)

Areal bone mineral density (aBMD) and bone mineral content (BMC) were measured for the whole body (without head, including the tail), as well as isolated to the lumbar spine (L3-L5) using a DXA X-ray cabinet system calibrated to bone density (UltraFocus^DXA^ , Hologic/Faxitron Bioptics, Tucson, AZ) as described previously(^21^). As detailed above, DXA images were acquired on the first day of PTH administration and at euthanasia.

### Mechanical testing via 3-point flexure

Fresh-frozen femurs were thawed and tested in 3-point flexure to measure whole bone structural mechanical properties as described previously (^22^). In short, femurs were placed anterior side down on 1 mm diameter supports spaced 10 mm apart, applying midspan loading to failure at 0.1 mm/second with force and displacement data acquired at 10 Hz (TA Instruments 3200, Eden Prairie, MN). Flexural strength was defined as the maximum applied force, with structural stiffness measured within the linear portion of the force-deflection response and work-to-fracture measured to the deflection at maximum force (strength). The distal half of the femur was then imaged via microCT to quantify indices of trabecular and cortical morphometry. Due to concerns of freeze-thaw processing of samples, we conducted studies to determine if mid-diaphyseal geometry must be imaged prior to mechanical tests. We determined that the diaphyseal geometry immediately adjacent (distal) to the midspan loading contact point (and fracture location) provided numerical measurements that were appropriate for evaluation of cortical bone, and equivalent to the numerical measures at the contact point, thus also providing geometric indices relevant to interpreting mechanical test results. Thus, this approach was incorporated into our phenotyping pipeline.

### Bone architecture

Trabecular bone morphometry was quantified in the distal metaphysis of the left femur and vertebral centrum of the L4 vertebra as described previously (^22–24^). Cortical bone morphometry was quantified at mid-diaphysis of the femur. Femurs and vertebrae were imaged in 70% ethanol using X-ray micro-computed tomography, collecting 2000 conebeam projections per revolution at 70 kVp (200μA) and an integration time of 500 msec within a 20 mm diameter field of view (μCT50, Scanco Medical AG, Bassersdorf, Switzerland). Three-dimensional images were reconstructed at 10 μm resolution using standard convolution back projection algorithms with Shepp and Logan filtering and rendered at a discrete voxel density of 1,000,000 voxels/mm^3^ (isometric 10 μm voxels). Bone mineral was calibrated to a discrete-step hydroxyapatite phantom and segmented from marrow and soft tissue in conjunction with a constrained Gaussian filter to reduce noise, applying mineral density thresholds of 550 and 700 mg HA/cm^3^ for trabecular and cortical bone, respectively. Volumetric regions for trabecular bone analysis were selected within the endosteal borders of the femoral metaphysis to include the secondary spongiosa located ∼1 mm (∼7% of length) from the growth plate and extending 5% of femur length proximally. Likewise, the central 80% of vertebral body height within the L4 vertebra was selected for trabecular analysis. Trabecular morphometric parameters were measured without imposing a presumed structural model to obtain direct measures of trabecular volume fraction (BV/TV), thickness (Tb.Th), number (Tb.N) and spacing (Tb.Sp). Cortical transverse morphometry was averaged within a mid-diaphyseal span of ∼5% of femur length to obtain measures of total cross-sectional area (bone + medullary area, mm^2^), cortical bone area (mm^2^), marrow area (mm^2^) and bone area fraction (%).

### Statistical analysis

All statistical analysis was conducted in R version 4.4.2. As there are known differences in bone biology between males and females, all analysis was conducted in a sex separated manner. Specific pair-wise comparisons between strains were not the goal of this study and therefore differences in PTH effects were determined by a two-sided unpaired T-Test within each strain. A multiple testing correction penalty was applied to the p values from these pair-wise comparisons using a Bonferroni method, and a p value less than 0.00625 was considered significant. For each phenotype of interest, an additive linear model (strain + treatment) was constructed and compared to an interactive model (strain * treatment). The difference between these models was tested by ANOVA to test for heterogeneity of effects across strains. A similar analysis was done to determine if there was a differential response to PTH between males and females.

To test for heritability, a simulated population of potential differences between the PTH treated animals and the saline treated animals within a strain and within a phenotype was created by conducting 10,000 bootstraps of the data, using the *boot* package for R (version 1.3-31). The *ICC* package for R (version 2.4.0) was used to calculate the unadjusted interclass correlation coefficient using strain as the classifier. This value was used to estimate the Broad Sense Heritability (H^2^) which is the portion of the phenotypic variance attributed to all types of genetic variation, as per the method suggested by Philip *et al* (^25^).

## RESULTS

### Bone mineral density and body composition

Administration of PTH did not affect body weight in males or females in any strain and had no effect on lean or fat mass in most strains (**Supplement Table 1**). As expected, intermittent exposure to PTH increased whole body BMD in B6 male and female mice, the genetic reference strain (**Figure 1, Supplement Table 2A&B**). A moderate BMD response was measured in both male and female NOD and PWK mice, but other strains showed sex-specific responses. Specifically, a BMD response was observed in male A/J, but not in female, and a moderate response was seen in female 129 mice but not in male (**Figure 1**). While the strain-by-treatment effects were not significant for either males or females, there was a strong difference when testing if males and females responded differently to PTH (p<0.001) for whole body BMD. The broad sense heritability for “BMD response to PTH treatment” was 85.0% for the females and 38.6% for the males (**Table 1**). This reflects the overall less robust BMD response to PTH observed in the males. The strain-specific response for whole-body bone mineral content (BMC) followed a similar pattern for the males, but only the B6 strain showed a significant response in the females (**Supplement Table 2A&B**). In the lumbar spine of male mice, only A/J moderately responded to PTH for BMD and BMC. In the lumbar spine of female mice, a positive response for BMD was detected in the CAST, NOD and PWK strains, and for BMC only in the B6 and PWK females (**Supplement Table 2A&B**). Baseline BMD data collected before treatment administration can be found in **Supplementary Table 2(C&D)**. It should be noted that the difference in the 12 week BMD for the female NZO is largely driven by one outlier value. This outlier was not removed as the p value as not significant after multiple testing correction. The percent difference (delta) between the pre-treatment BMD and the post-treatment BMD is also provided, but require caution for statistical interpretation. A delta analysis compounds the measurement error 1.41-fold as it incorporates two tests, resulting in a larger standard deviation and requires increasing the multiple testing correction penalty to 0.003.

**Figure 1.**
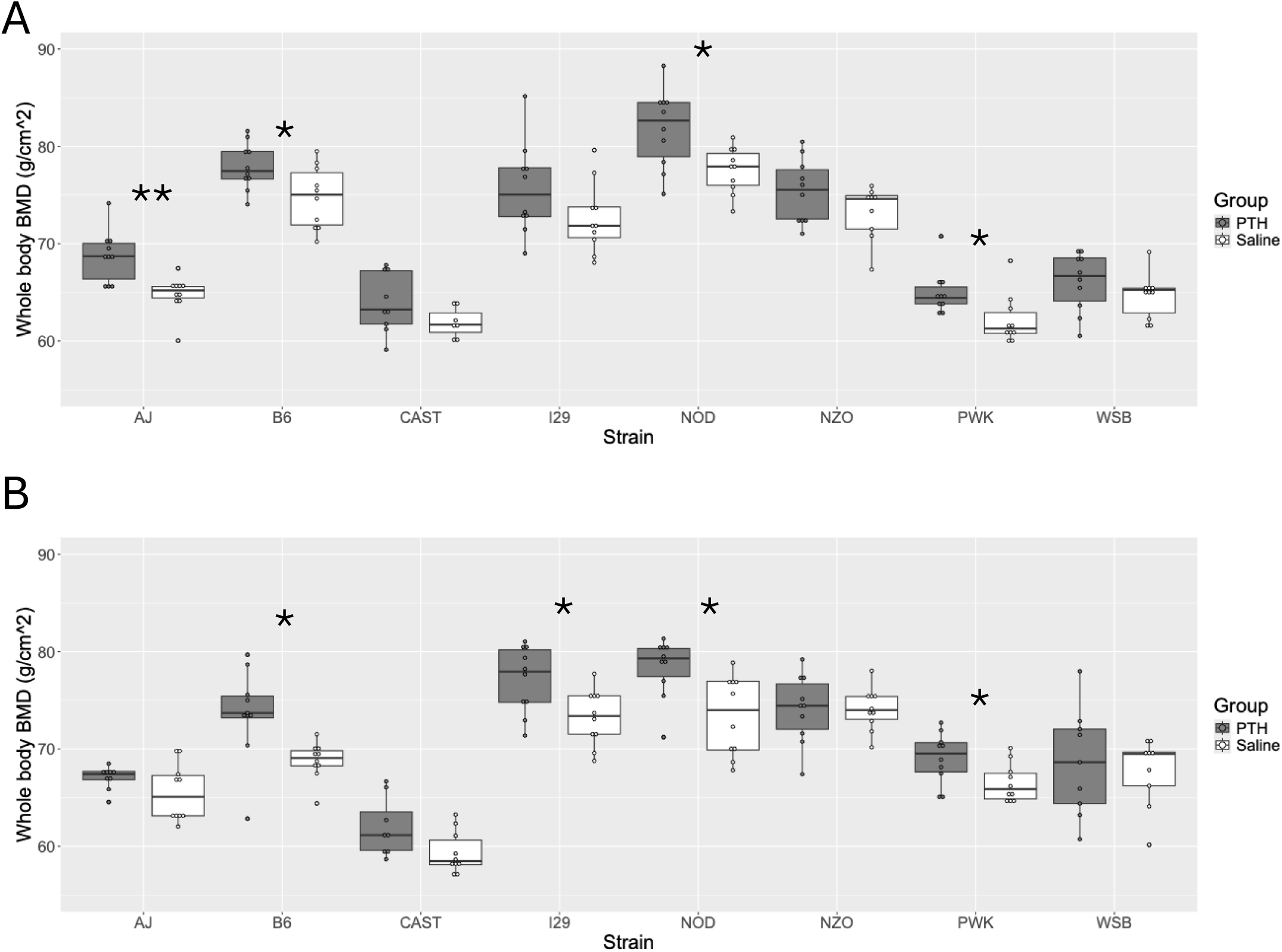
**<u>Whole body BMD response to intermittent PTH</u>**. Whole body bone mineral density (BMD) sans the head is shown for male (A) and female (B) mice of 8 different inbred strains after 4 weeks of treatment. PTH treated animals are shown in grey and saline treated animals in white. Values for PTH treated animals are shown in grey and saline treated animals in white. **p< 0.00625, *p<0.05. The value of p< 0.00625 was chosen for significance after Bonferroni multiple testing correction. Complete P values and per strain means can be found in Supplemental materials. For the interaction between strain and treatment, p=0.83 for the males and p=0.1605 for the females indicating that strain specific responses to treatment were not significant.

**Table 1.** Broad sense heritablities (H^2^, %) for response to PTH.

| **Phenotype** | **Male** | **Female** |
| --- | --- | --- |
| BMD | 38.9 | 85.0 |
| Cortical Area | 71.0 | 39.1 |
| Breaking Strength | 56.0 | 49.7 |
| BV/TV (femoral) | 64.8 | 87.8 |
| BV/TV (vertebral) | 77.2 | 76.1 |

**Table 2.** Summary of the impact of PTH by sex, bone compartment and anatomic location.

| Strain | Sex | Whole body BMD | Cortical Area | Breaking Strength | Femoral BV/TV | Vertebral BV/TV |
| --- | --- | --- | --- | --- | --- | --- |
| 129 | Male |  |  |  |  |  |
|  | Female | ↑ | ↑ |  | ↑ |  |
| A/J | Male | ↑ | ↑ | ↑ | ↑ | ↑ |
|  | Female |  |  |  |  |  |
| B6 | Male | ↑ |  |  |  |  |
|  | Female | ↑ | ↑ | ↑ |  |  |
| CAST | Male |  | ↑ |  | ↓ |  |
|  | Female |  |  |  | ↓ |  |
| NOD | Male | ↑ |  |  | ↓ | ↑ |
|  | Female | ↑ | ↑ | ↑ |  | ↑ |
| NZO | Male |  |  |  |  | ↑ |
|  | Female |  |  |  |  |  |
| PWK | Male | ↑ | ↑ | ↑ |  | ↑ |
|  | Female | ↑ | ↑ |  |  | ↑ |
| WSB | Male |  |  |  |  |  |
|  | Female |  |  |  | ↑ | ↑ |

### Femoral Cortical Architecture and strength

In males, we observed a moderate increase with PTH treatment for cortical area in A/J and CAST mice and a larger response was seen in PWK mice. A change in intramedullary area was detected only in the CAST strain. In females, only NOD mice were observed to have an increase in cortical area (**Figure 2, Supplemental Table 3 A&B**). Change in cortical area in response to PTH moderately was highly heritable (H^2^ = 71%, **Table 1**) in males and moderately so in females at 39.1% (**Table 1**). There was a strong effect of sex on response to treatment for cortical area (p<0.0001) and a more moderate effect for femur strength (p=0.001). No change in intramedullary area in the males was observed. The A/J and CAST female mice experienced a decrease in intramedullary area and an increase in cortical thickness, but no significant change in cortical area. No change in periosteal circumference was noted for either strain (A/J, PTH = 4.38±0.204 vs. Saline = 4.74±0.244 mm, p = 0.999; CAST, PTH = 3.88±0.193 vs. Saline = 4.01±0.216 mm, p = 0.910).

**Figure 2.**
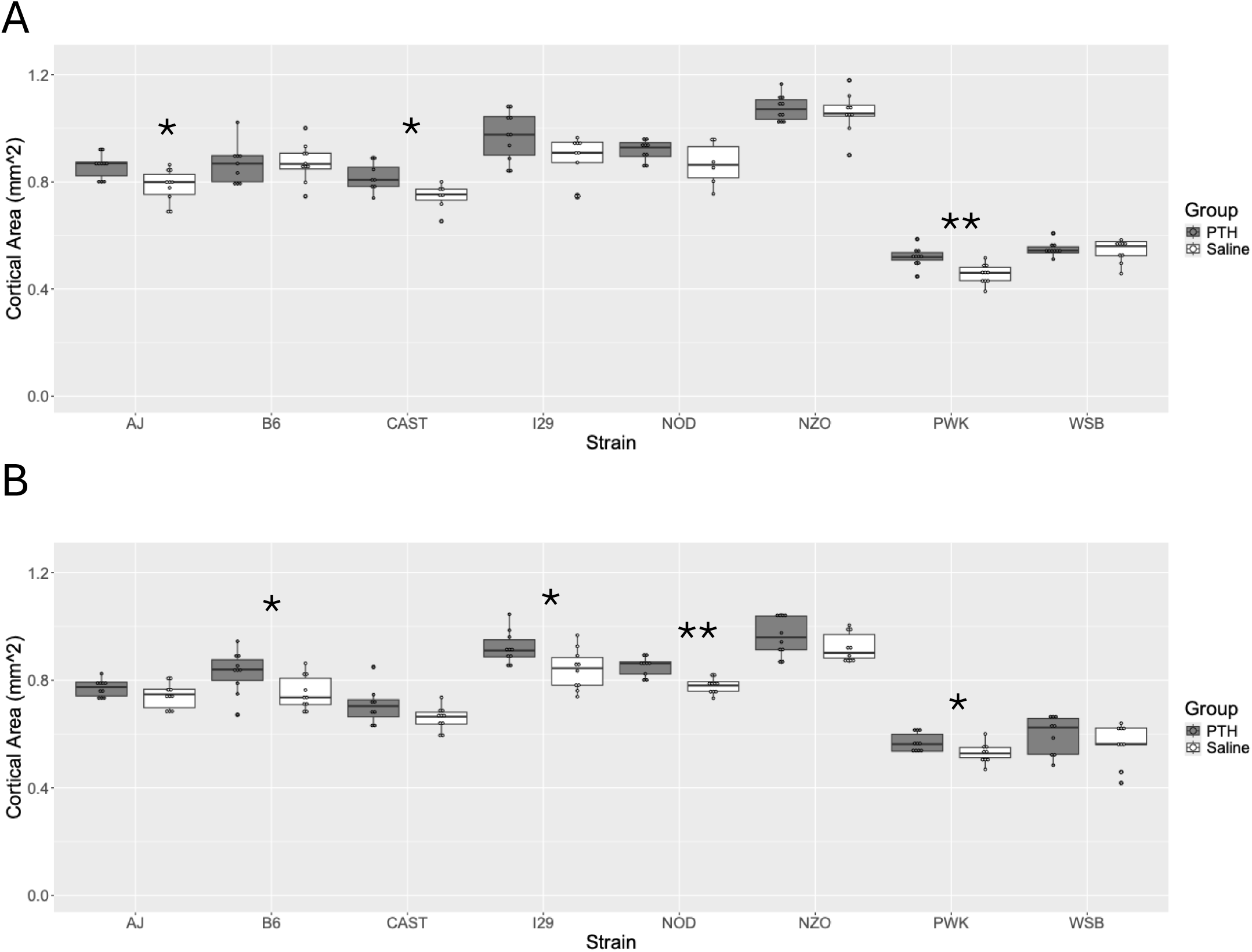
**<u>Femoral cortical area response to intermittent PTH</u>**. Cortical geometry at the mid diaphysis was measured by μCT after 4 weeks of PTH treatment. Cortical area in PTH treated (grey) and saline treated (white) males (A) and females (B) is present. **p< 0.00625, *p<0.05. The value of p< 0.00625 was chosen for significance after Bonferroni multiple testing correction. Complete P values and per strain means can be found in Supplemental materials. For the interaction between strain and treatment, p=0.168 for the males and p=0.707 for the females indicating that strain specific responses to treatment were not the whole significant.

Increases in femur strength with PTH treatment were seen in male A/J and PWK, and in female B6 mice (**Figure 3, Supplemental Table 4 A&B**). The heritability for change in breaking strength after PTH treatment was moderate for both males and females (**Table 1**). This may reflect the high variance in this phenotype. For all groups showing an increase in breaking steength, there was a concordant increase in cortical area. A change (increase) in flexural stiffness was observed only in the NOD males, which did not exhibit a change in breaking strength with PTH treatment, and in B6 females, which demonstrated concomitant increase in strength. PTH did not affect work-to-failure in any of the groups. Flexural and torsional second moments of area, which are geometrical measures of strength and stiffness, largely followed the pattern of changes in cortical area.

**Figure 3.**
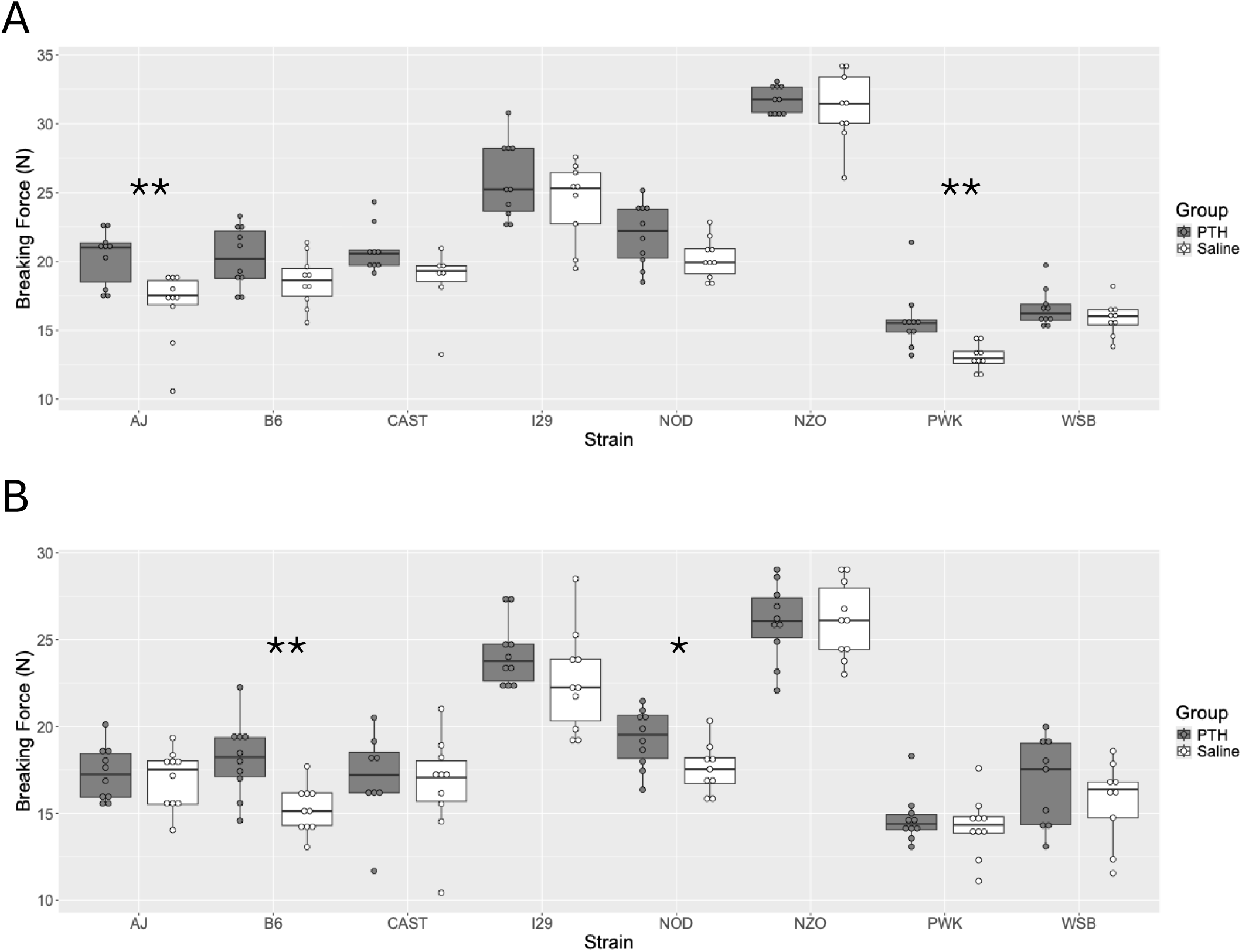
**<u>Femoral Breaking Strength in saline and PTH treated mice</u>**. Femoral breaking strength was measured by 3-point bending in female (A) and male (B) of 8 inbred strains. Values for PTH treated animals are shown in grey and saline treated animals in white. **p< 0.00625, *p<0.05. The value of p< 0.00625 was chosen for significance after Bonferroni multiple testing correction. Complete P values and per strain means can be found in Supplemental materials. For the interaction between strain and treatment, p=0.403 for the males and p=0.358 for the females indicating that strain specific responses to treatment were not the whole significant.

### Femoral and vertebral trabecular bone

In male mice we observed a decrease in femur distal metaphysis trabecular bone volume fraction (BV/TV) in CAST and NOD strains (**Figure 4A and Supplemental Table 5A)**. In the A/J mice, there was a nominal increase in femoral trabecular volume fraction, coincident with thicker trabeculae and a large increase in trabecular number. In the CAST males, there was a trend towards thinner trabeculae with PTH treatment, but valid measurements of trabecular number could not be performed for the very sparse trabecular volume (**Figure 4**). The morphometric change driving the decrease could therefore not be determined. In the NOD males, this decrease in trabecular volume was associated with a reduction in the number of trabeculae, but unlike in the CAST males, the NOD males experienced no change in trabecular thickness. In the B6 and NZO males, we observed a large increase in trabecular thickness that was coincident with a decrease in trabecular number. The result was no net change in femur trabecular volume fraction. Both the response to treatment by strain (**Figure 4)** and the response to treatment by sex (p<0.001) were highly significant for femoral trabecular volume fraction.

**Figure 4.**
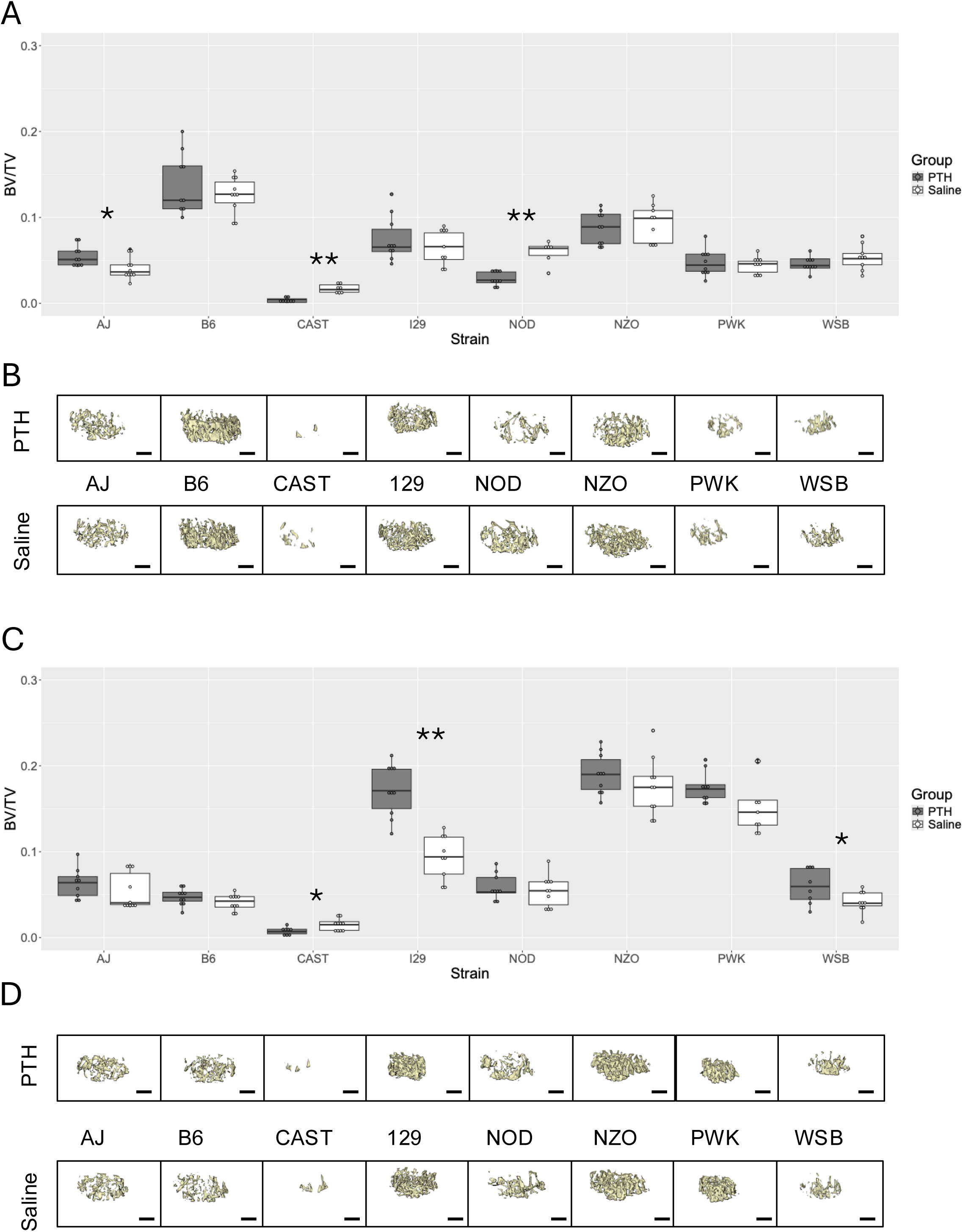
**<u>Distal femoral trabecular bone volume fraction (BV/TV) in saline and PTH</u> <u>treated mice</u>**. Distal femoral volume fraction (BV/TV) was measured by μCT after 4 weeks of PTH treatment. Date from the PTH treated (grey) and saline treated (white) males (A) and females (C) is present. Representative reconstructions of the trabecular compartment are shown for males (B) and females (D) such that the PTH treated animals are on the top row and the saline treated are on the bottom row. **p< 0.00625, *p<0.05. The value of p< 0.00625 was chosen for significance after Bonferroni multiple testing correction. Complete P values and per strain means can be found in Supplemental materials. For the interaction between strain and treatment, p < 0.0001 for both the males and the females indicating that strain specific responses to treatment were true.

In the strains that experienced a response in vertebral trabecular bone volume (BV/TV), change was always in the positive direction in the males (**Figure 5A & B and Supplemental Table 6A)**. As occurred in the distal femur, a response was measured in the L4 vertebrae in both the A/J and NOD males. A substantial increase in BV/TV with PTH treatment was measured in the L4 vertebra of NZO and PWK males as well. For all four strains, this increase in BV/TV corresponded to an increase in trabecular thickness, but unlike in the distal femur, change in trabecular number was not associated with this increase. As was observed in the femur, trabecular thickness increased in the vertebral body, but trabecular number decreased in the B6 males, with no net change in BV/TV. There was sufficient trabecular bone in the vertebral body in the CAST males to measure trabecular number. In response to PTH, trabecular number decreased substantially in this strain, but the decrease was not associated with any change in BV/TV.

**Figure 5.**
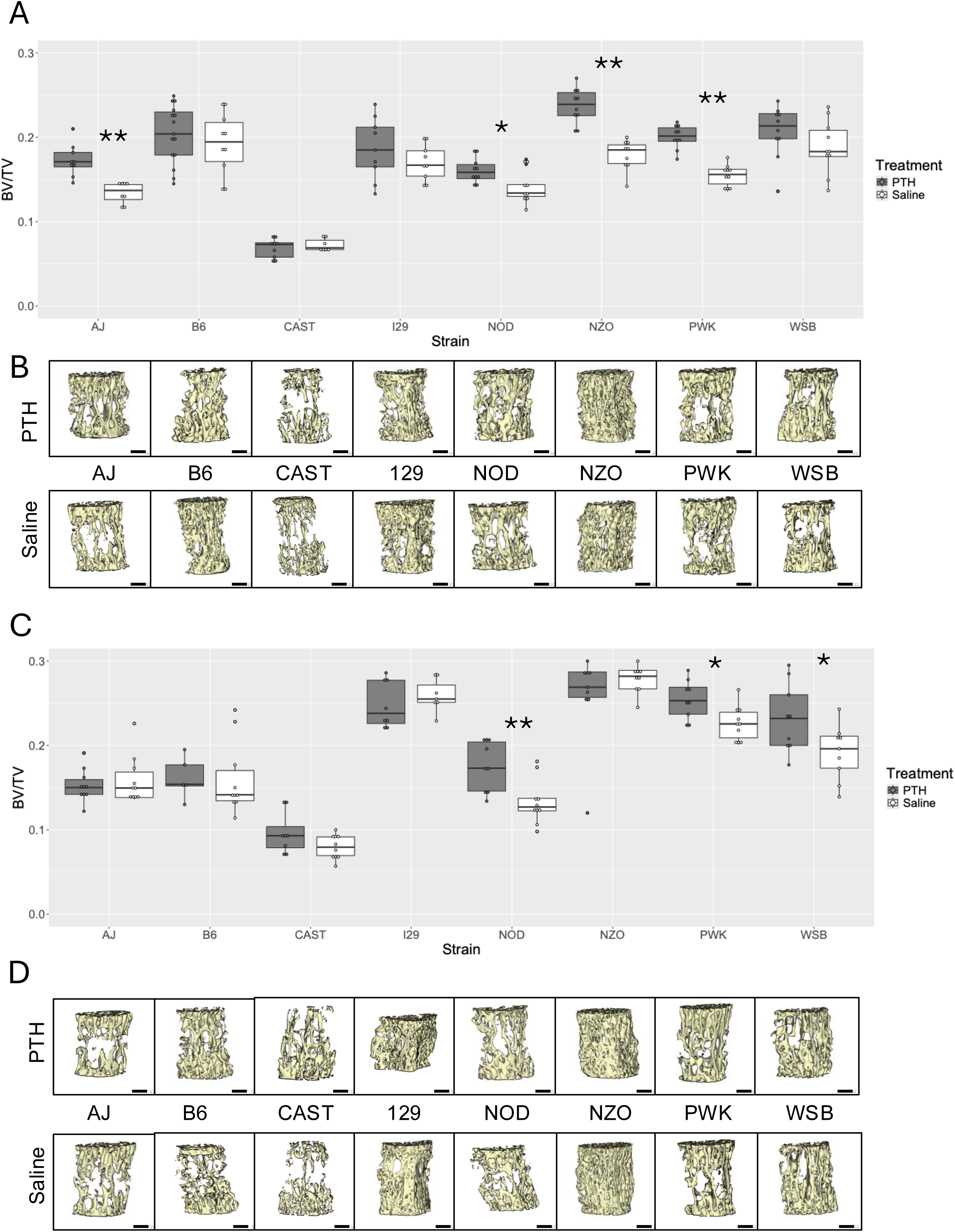
**<u>Lumbar vertebral trabecular bone volume fraction (BV/TV) in saline and PTH</u> <u>treated mice</u>**. Volume fraction (BV/TV) of the fourth lumbar vertebrae was measured by μCT after 4 weeks of PTH treatment. Date from the PTH treated (grey) and saline treated (white) males (A) and females (C) is present. Representative reconstructions of the trabecular compartment are shown for males (B) and females (D) such that the PTH treated animals are on the top row and the saline treated are on the bottom row. **p< 0.00625, *p<0.05. The value of p< 0.00625 was chosen for significance after Bonferroni multiple testing correction. Complete P values and per strain means can be found in Supplemental materials. For the interaction between strain and treatment, p=0.0003 for the males and p=0.0005 for the females indicating that strain specific responses to treatment were true.

A very different group of strains presented with a change in BV/TV after PTH treatment in the females, compared to the males, for the distal femur (**Figure 4C & D and Supplemental Table 5B)**. Here, BV/TV increased in the 129s. This was associated with an increase in trabecular thickness and no change in trabecular number. Conversely, in the WSB, where there was modest increase in BV/TV, there was no change in trabecular thickness but there was a dramatic increase in trabecular number. However, as occurred for the males, CAST females experienced a decrease in distal femoral BV/TV with PTH treatment. Unlike in the males, this was not associated with any change in trabecular thickness, and trabecular number could not be measured. In the vertebral body BV/TV increased in the NOD females treated with PTH (**Figure 5C & D and Supplemental Table 6B)**. This increase was accomplished by increasing trabecular thickness, with no associated change in trabecular number. In the WSB females, PTH treatment did not result in a statistically significant change in either trabecular thickness or trabecular number despite the substantial increase in BV/TV. Overall, the response to PTH was most heritable for trabecular volume fraction (BV/TV) at both the distal femur and in the vertebral body (**Table 1)**. As for the femur, the response to treatment for BV/TV by sex was highly significant (p<0.001) for vertebral BV/TV.

## DISCUSSION

The eight strains of mice used in this study were selected as they are the founder strains for a very powerful genetic mapping population called the Diversity Outbred(^14^). This resource is increasingly becoming the preferred outbred mouse model for genetic mapping studies in mice. The genomes of these 8 strains have been sequenced and among these strains there are ∼37.8 million single nucleotide polymorphism (SNP) differences and ∼6.1 million insertion/deletions (INDELs) (^26^). In comparison, the 1000 genomes project has suggested that only 2.7 million SNPs appear in humans with a minor allele frequency of greater than 1% (^27^). These eight strains represent a considerable amount of genetic variability, allowing for the estimation of heritability of the response to teriparatide/PTH in a relatively small number of mice. There are many advantages to studying drug response heritability in mice compared to humans. In mice, the environment is tightly controlled, including temperature exposure, light/dark cycle, and diet. There are no concerns for complicating factors such as medication use or differences in activity levels. Moreover, all mice within an inbred strain can be considered genetic clones of all other mice of that strain (^16^), allowing for repeated measures of phenotypes to overcome subtle differences that could be masked due to technical or measurement error. Further, by using mice we can collect data that cannot be collected in human cohorts such as bone strength and stiffness.

As more data is accumulated, it is becoming clearer that treatment failure and inadequate clinical response after teriparatide therapy is a significant problem for patients. A retrospective analysis of clinical trials data has shown that dose and duration of treatment matter when considering changes in BMD (^6^). A limitation of that analysis was that the clinical trial data were available only for women. At 12 months of dosing with 20ug/day, the retrospective analysis showed that 13% of women did not attain a 3% or more increase in BMD at the lumbar spine and 68% did not attain a similar increase in BMD at the hip. At 18 months, the non-responder rate based on BMD ranged from 6 to 9% for the lumbar spine and 50 to 53% for the hip(^6^). These clinical trial data highlight the magnitude of the problem of non-response in patients. Unlike for many other drugs(^28^), we have no pharmacogenomics testing that can *a priori* predict response to teriparatide. In our study, half of the strains responded in both males and females for whole body BMD, but there were differences in which strains responded between the two sexes. Our data strongly suggest that response to PTH is genetic and sex specific, but a study in outbred mice will be needed to test more complex allelic distributions and to determine the genes responsible for such a genetic response. As in humans, we found that some mice were very resistant to PTH. WSB males did not respond for any of the phenotypes we measured and A/J females only showed a response for the phenotypes of intramedullary area and cortical thickness.

Fracture after treatment with teriparatide remains an issue for some patients. In a retrospective study of men and women, Elraiyah et al found that up to 7% of patients continued to fracture after at least 6 months of teriparatide treatment(^8^). In this cohort of men and women, 34.8% of patients failed to achieve a 3% or greater increase in BMD at the spine, hip, or both sites. In a retrospective study of osteoporotic treatment effectiveness in post-menopausal women (the TAILOR study), an 11% treatment failure rate was observed in patients treated with teriparatide (^29^). In that study, failure was defined as two or more incident fractures after the initiation of treatment (^30^). In our mouse study, the heritability for increased femur flexural strength after PTH treatment was 56% in males and 49.2% females (**Table 1**), indicating that half of the variation in response to PTH for femur strength can be attributed to mouse strain/genetics. Based on our mouse femur data, we postulated that just as occurred for changes in BMD (^11^), fracture after teriparatide treatment can in part be explained by a patient’s genetics. It is not practical to collect such data in humans, as heritability of fracture is very difficult to estimate, and the data do not exist in large enough numbers to provide for matched treated and untreated cohorts for fracture (^31,32^). Our data showed that increased femur strength at mid-diaphysis coincided with an increase in the amount of cortical bone. The mechanism by which bone strength is changed is very different in mice versus humans, as teriparatide treatment was not associated with an increase in cortical thickness in either pre-menopausal (^33^) or post-menopausal women (^34^).

A general practice for studies investigating teriparatide response has been to compare studies in men to literature reported data collected in women as little data has been collected for men (^35^). A single retrospective study of men and women that had completed at least 12 months of teriparatide therapy, Niimi *et al* showed that there was no difference between men and women for the absolute increase in lumbar spine BMD and femoral neck BMD, but longitudinally their data suggested that women were more responsive than men at the femoral neck at earlier timepoints (^36^). Our study was able to directly compare males and females within and between different genetic backgrounds, highlighting important sex-specific responses. For example, in all strains that showed a cortical response to PTH, treatment induced an increase in cortical area but the mechanism by which this happened was different among the strains and by sex. Regardless of sex, the increase in cortical area was largely associated with an increase in cortical thickness. In male mice, intramedullary area did not change except in the CAST mice. This suggests that in male mice cortical bone mass was most commonly increased by periosteal expansion. In female mice, a decrease in intramedullary area was more common, suggesting a trend towards endosteal expansion instead. These mice were 12 weeks of age at the initiation of treatment. By definition, all of these mice could be considered sexually mature, but mice at this age are still actively growing (^37^). This male/female difference in periosteal expansion is highly reminiscent of that seen in humans surrounding puberty and with aging (^38^). There were clear deviations to this trend in select strains. Specifically, CAST males and females presented with decreased intramedullary area and neither PWK males nor females showed any significant changes in intramedullary area. This suggests that while there are sex-specific patterns, genetic factors may be able to override them.

Multiple clinical studies have shown that the skeletal anabolic response to intermittent teriparatide is stronger at the spine than at the hip (^6^). This is reflected in fracture data which show that regardless of dose, teriparatide is more effective at reducing vertebral fractures than non-vertebral fractures(^39^). In randomized control studies the overlap between non-responders at the spine and hip was not perfect (^6^). In other words, some patients responded at both sites whereas some patients responded at the hip but not the spine and vice versa. We saw a similar phenomenon in our data, but our data can provide more granularity within the cortical and trabecular compartments of different anatomic sites. For the males, only one strain (NOD) showed a change in trabecular bone volume (BV/TV) in both lumbar vertebrae and the distal femur. In females, there was no overlap in response to PTH for trabecular bone in the lumbar spine and femur among the strains that responded to treatment. This is supported by studies showing that osteoblast gene expression and mineralizing ability is different for cells taken from different anatomic sites (^40^).

We saw a decrease in BV/TV in the femur in male B6, CAST and NOD mice and in CAST female mice. A similar phenomenon was not seen in the vertebra. Studies of single doses of teriparatide showed that resorption is immediately increased after dosing, before formation begins to increase (^41^). The kinetics of turnover in different mouse strains is poorly understood, but it is known that osteoblast mediated formation and osteoclast number are both independently genetically controlled (^42,43^). A limitation of this study is that it was not possible to measure serum bone turnover markers in all strains of mice as the wild derived mice are very small, precluding safe blood draws over time. We hypothesize, based on known differences in inbred strains and transgenic mouse models, that some strains were more sensitive to the PTH with regards to ramping up bone resorption in the femur. This same reaction to teriparatide, however, may happen in humans. Using HR-pQCT, a decrease in BV/TV was observed at the distal radius and the distal tibia in women after 18 months of teriparatide therapy. The authors of this study speculate that teriparatide resulted in an increase in number of trabeculae, albeit each one smaller and thinner in size (^44^), but this is not what we observed in mice. In the male mice that experienced a decrease in BV/TV, trabecular number also decreased. We did observe that an increase in trabecular number could be associated with an increase in osteoclast number in some transgenic mice (^42^).

## CONCLUSIONS

As this study of mice has repeated measures within a given genotype, we can conclusively show that there are non-responders to PTH treatment, but our data clearly show that response is heavily driven by genetic factors. Further, these data suggest that a primary driver of skeletal response to PTH stems from interactions between genetics and sex. As this is a mouse study, future work will need to be conducted to determine if similar sex by genetic interactive effects are seen in humans. This work supports the future use of outbred mice to conduct GWAS studies for the response of bone to intermittent PTH to determine the genes responsible for mediating the effect of intermittent PTH on bone.

## ACKNOWLEDGEMENTS

The authors would like to thank Michael David, PhD for his assistance with analyzing the μCT data. The authors would like to thank Dr. C. Gignoux for his advice on calculating heritabilities. The graphical abstract was generated using BioRender.

## FUNDING

This work was funded by the following grant to DJA and CLAB from NIH/NIAMS: AR070879. The stipend for SER was supported by the Pathways in Genomic Data Science program funded through the National Human Genome Research Institute (R25HG012919)

## DATA AVAILABLITY

All raw phenotyping data is available upon request. No specialized code was generated for the data analysis, and all analysis was conducted using the open-source packages for R listed in the methods.

## SUPPLEMENTARY MATERIAL

Supplemental Table 1: Body weight and composition for males (A) and females (B).

Supplemental Table 2: Whole body BMD and BMC for males (A) and females (B) as measured by DXA at 16 weeks and BMD measured at 12 weeks (baseline, C&D).

Supplemental Table 3: Femoral cortical architecture as measured by μCT for males (A) and females (B)

Supplemental Table 4: Femoral cortical mechanical properties as measured by 3-point bending for males (A) and females (B)

Supplemental Table 5: Femoral distal trabecular architecture as measured by μCT for males (A) and females (B)

Supplemental Table 6: Vertebral (L4) trabecular architecture as measured by μCT for males (A) and females (B)

## AUTHOR CONTRIBUTIONS

DJA: Obtained funding for this work and conceptualized the hypothesis. Contributed to data analysis, figure construction and writing of the final manuscript.

DAG: Designed the experimental protocol. Collected most of the raw data. Contributed to data analysis.

SER: Contributed to data analysis and statistical method development.

RDM: Helped with experimental design and execution. Did critical reviews of the final manuscript.

NSS: Developed methods for high throughput testing of bone strength in diversely shaped specimens. Contributed to data analysis.

ELM: Advised on the statistical analysis plan and aided in interpretation of said analysis.

CLAB: Obtained funding for this work and conceptualized the hypothesis. Assisted with experimental design. Conducted to data analysis, statistical analysis, figure construction and writing of the final manuscript.

Raw animal phenotyping data is available upon request and/or is provided in the supplemental materials.

This work was funded by NIH/NIAMS AR073346 awarded to CLAB and DJA. This work was supported by Pathways in Genomic Data Science through the National Human Genome Research Institute (R25HG012919).

All use of animals was approved by the Institutional Animal Care and Use Committee (IACUC) of the University of Colorado (CU), Anschutz Medical Campus (Protocol number: 00902).

This study does not include data from human subjects.

---

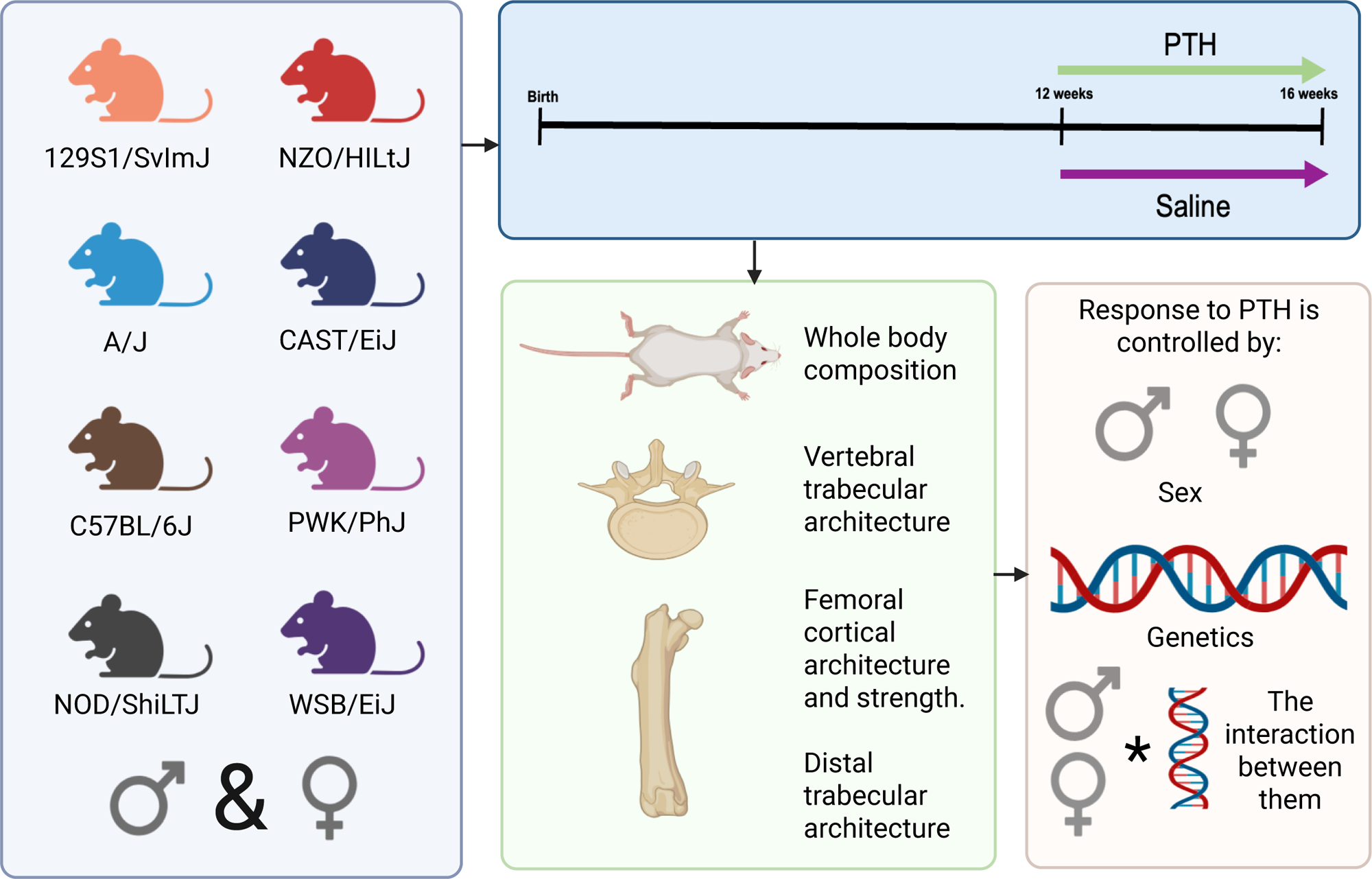

## Notes

### Competing Interest Statement

The authors have declared no competing interest.

### Summary of Updates

This version has been updated to reflect reviewers' requests for revision. This includes adding more data to the supplement and providing a clearer explanation of the multiple-testing correction conducted. The title has been changed to include the word mouse.

