## Supplementary material for "Bone response to intermittent parathyroid hormone (PTH) is both genetic and sex specific in mice": Supplmental tables

**Supplemental Table 1**. Body weight and composition

A. **Male**

|  |  | Body Weight (g) | | Lean Mass (g) | | Fat Mass (%) | |
| --- | --- | --- | --- | --- | --- | --- | --- |
| Strain | Treatment  (N) | Average | P value | Average | P value | Average | P value |
| 129 | Saline (10) | 26.38±1.66 | 0.3370 | 18.50±1.47 | 0.3173 | 21.80±4.34 | 0.3978 |
|  | PTH (10) | 27.15±1.95 |  | 19.16±1.42 |  | 20.15±4.17 |  |
| A/J | Saline (10) | 26.07±1.95 | 0.8459 | 16.26±0.93 | 0.4445 | 28.01±3.24 | 0.1742 |
|  | PTH (10) | 25.9±1.91 |  | 16.61±1.05 |  | 26.31±1.87 |  |
| B6 | Saline (10) | 28.4±1.93 | 0.8257 | 20.39±1.26 | 0.6924 | 21.18±3.13 | 0.5269 |
|  | PTH (10) | 28.59±1.87 |  | 20.10±1.85 |  | 22.17±3.67 |  |
| CAST | Saline (7) | 17.36±0.64 | 0.6253 | 13.27±0.90 | 0.6931 | 15.58±2.78 | 0.8889 |
|  | PTH (9) | 17.1±1.36 |  | 13.08±0.94 |  | 15.8±3.57 |  |
| NOD | Saline (10) | 29.42±1.78 | 0.0854 | 21.60±1.58 | 0.2292 | 17.88±3.43 | 0.6343 |
|  | PTH (10) | 30.74±1.44 |  | 22.49±1.62 |  | 17.17±3.18 |  |
| NZO | Saline (9) | 50.96±4.87 | 0.7113 | 29.64±1.87 | 0.8971 | 34.28±5.33 | 0.1807 |
|  | PTH (10) | 50.28±2.96 |  | 29.79±3.00 |  | 30.96±6.18 |  |
| PWK | Saline (10) | 17.34±1.15 | 0.0587 | 13.55±0.88 | 0.1911 | 11.84±4.33 | 0.0561 |
|  | PTH (10) | 18.23±0.76 |  | 14.04±0.71 |  | 8.12±3.79 |  |
| WSB | Saline (10) | 16.91±1.27 | 0.8444 | 14.28±1.07 | 0.4668 | 6.13±1.99 | 0.0637 |
|  | PTH (10) | 16.81±0.88 |  | 13.96±0.82 |  | 8.05±2.34 |  |

* p <0.05, bold: p<0.00625

B. **Female**

|  |  | Body Weight (g) | | Lean Mass (g) | | Fat Mass (%) | |
| --- | --- | --- | --- | --- | --- | --- | --- |
| Strain | Treatment  (N) | Average | P value | Average | P value | Average | P value |
| 129 | Saline (10) | 21.46±1.69 | 0.4569 | 15.04±0.78 | 0.6928 | 16.89±2.64 | 0.2666 |
|  | PTH (10) | 20.97±1.13 |  | 14.19±0.65 |  | 15.72±1.84 |  |
| A/J | Saline (9) | 21.09±1.23 | 0.1685 | 14.71±0.74 | 0.5215 | 23.09±4.11 | 0.1255 |
|  | PTH (10) | 20.3±1.23 |  | 13.97±0.63 |  | 20.52±2.76 |  |
| B6 | Saline (10) | 20.70±1.59 | 0.4509 | 14.37±0.92 | 0.2056 | 22.38±3.46 | 0.3234 |
|  | PTH (10) | 21.21±1.36 |  | 15.10±1.49 |  | 20.19±5.85 |  |
| CAST | Saline (8) | 14.10±1.54 | 0.1981 | 10.81±0.94 | 0.2994 | 11.89±4.34 | 0.0299* |
|  | PTH (10) | 13.30±1.03 |  | 10.36±0.85 |  | 7.80±2.89 |  |
| NOD | Saline (10) | 21.83±1.37 | 0.9618 | 16.25±1.41 | 0.9972 | 14.31±3.41 | 0.6416 |
|  | PTH (10) | 21.80±1.39 |  | 16.25±1.08 |  | 13.59±3.39 |  |
| NZO | Saline (10) | 43.19±3.37 | 0.9960 | 24.00±1.12 | 0.0094* | 39.49±5.30 | 0.2724 |
|  | PTH (10) | 43.20±5.22 |  | 22.66±0.95 |  | 39.85±7.70 |  |
| PWK | Saline (10) | 17.08±1.36 | 0.1758 | 12.97±0.88 | 0.7736 | 10.61±3.91 | 0.8715 |
|  | PTH (10) | 16.13±1.64 |  | 12.85±1.00 |  | 10.91±4.38 |  |
| WSB | Saline (9) | 16.59±2.30 | 0.6432 | 12.74±1.15 | 0.3352 | 8.13±1.97 | 0.4424 |
|  | PTH (9) | 16.09±2.19 |  | 13.27±1.09 |  | 7.25±2.69 |  |

* p <0.05, bold: p<0.00625

**Supplemental Table 2**. Areal whole body and lumbar BMD and BMC.

A. **Male – 16 wks**

|  |  | Whole body | | | | Lumbar Spine | | | |
| --- | --- | --- | --- | --- | --- | --- | --- | --- | --- |
|  |  | BMD (mg/cm^2^) | | BMC (mg) | | BMD (mg/cm^2^) | | BMC (mg) | |
| Strain | Treatment  (N) | Average | P value | Average | P value | Average | P value | Average | P value |
| 129 | Saline (10) | 72.6±3.62 | 0.1255 | 604.8±38.6 | 0.1339 | 81.3±6.51 | 0.5700 | 33.0±2.79 | 0.5537 |
|  | PTH (10) | 75.6±4.69 |  | 635.8±48.8 |  | 79.6±6.30 |  | 33.8±3.46 |  |
| A/J | Saline (10) | 64.8±1.93 | **0.0016** | 476.9±29.3 | **0.0022** | 73.6±6.22 | 0.0318* | 26.3±2.64 | 0.0271* |
|  | PTH (10) | 68.7±2.67 |  | 512.4±26.3 |  | 80.3±6.7 |  | 29.5±3.18 |  |
| B6 | Saline (10) | 74.8±3.19 | 0.0223* | 566.1±40.8 | 0.0729 | 83.9±7.44 | 0.2902 | 32.3±4.01 | 0.7927 |
|  | PTH (10) | 77.9±2.40 |  | 598.8±37.4 |  | 87.1±5.54 |  | 32.7±3.63 |  |
| CAST | Saline (7) | 61.9±1.55 | 0.1142 | 354.7±10.9 | 0.4973 | 59.2±3.69 | 0.5053 | 16.3±1.72 | 0.9314 |
|  | PTH (9) | 63.9±3.08 |  | 362.5±31.1 |  | 61.0±6.87 |  | 16.4±2.70 |  |
| NOD | Saline (10) | 77.5±2.37 | 0.0111* | 623.9±25.6 | 0.0417* | 87.8±4.69 | 0.2269 | 33.7±2.37 | 0.8869 |
|  | PTH (10) | 81.8±4.04 |  | 659.2±43.0 |  | 91.3±7.53 |  | 33.6±3.09 |  |
| NZO | Saline (9) | 73.2±2.78 | 0.1295 | 933.1±89.0 | 0.9947 | 82.1±7.03 | 0.3968 | 49.1±5.27 | 0.5118 |
|  | PTH (10) | 75.4±3.28 |  | 933.4±85.5 |  | 84.7±5.96 |  | 47.7±3.87 |  |
| PWK | Saline (10) | 62.2±2.53 | 0.0178* | 355.3±24.0 | **0.0023** | 61.7±4.30 | 0.0711 | 17.2±2.50 | 0.0842 |
|  | PTH (10) | 65.0±2.31 |  | 390.0±19.2 |  | 65.1±5.60 |  | 18.8±0.84 |  |
| WSB | Saline (10) | 64.6±2.9 | 0.2493 | 391.5±28.5 | 0.5303 | 65.3±5.79 | 0.5655 | 17.6±2.73 | 0.7857 |
|  | PTH (10) | 66.1±3.04 |  | 400.4±33.6 |  | 66.8±5.60 |  | 17.9±2.40 |  |

* p <0.05, bold: p<0.00625

B. **Female – 16 wks**

|  |  | Whole body | | | | Lumbar Spine | | | |
| --- | --- | --- | --- | --- | --- | --- | --- | --- | --- |
|  |  | BMD (mg/cm^2^) | | BMC (mg) | | BMD (mg/cm^2^) | | BMC (mg) | |
| Strain | Treatment  (N) | Average | P value | Average | P value | Average | P value | Average | P value |
| 129 | Saline (10) | 73.2±2.88 | 0.0132* | 576.8±51.2 | 0.4349 | 90.3±5.74 | 0.8090 | 35.3±5.33 | 0.9425 |
|  | PTH (10) | 77.1±3.41 |  | 592.0±30.6 |  | 89.5±8.45 |  | 31.1±4.06 |  |
| A/J | Saline (9) | 65.5±2.95 | 0.1624 | 452.4±36.6 | 0.9419 | 79.9±8.74 | 0.3822 | 26.7±5.62 | 0.1748 |
|  | PTH (10) | 67.0±1.19 |  | 451.3±26.5 |  | 77.1±3.62 |  | 24.1±2.05 |  |
| B6 | Saline (10) | 68.8±1.91 | 0.0107* | 437.1±28.8 | **0.0035** | 80.9±6.22 | 0.2629 | 27.7±2.28 | **0.0051** |
|  | PTH (10) | 73.6±4.65 |  | 528.5±42.4 |  | 84.8±8.52 |  | 33.4±2.79 |  |
| CAST | Saline (8) | 59.3±2.15 | 0.0656 | 313.4±20.9 | 0.3588 | 57.2±4.74 | 0.0487* | 14.6±1.43 | 0.2333 |
|  | PTH (10) | 61.9±3.03 |  | 324.7±27.6 |  | 61.8±4.21 |  | 15.6±2.01 |  |
| NOD | Saline (10) | 73.4±4.09 | 0.0072* | 551.9±32.4 | 0.0074* | 85.5±7.78 | 0.0207* | 30.3±2.72 | 0.0679 |
|  | PTH (10) | 78.4±3.07 |  | 591.1±24.5 |  | 93.3±5.78 |  | 32.7±2.78 |  |
| NZO | Saline (10) | 74.1±2.19 | 0.9830 | 844.1±59.0 | 0.1809 | 89.3±7.06 | 0.5828 | 50.9±5.33 | 0.5382 |
|  | PTH (10) | 74.1±3.51 |  | 905.6±124.2 |  | 87.4±8.35 |  | 52.5±5.92 |  |
| PWK | Saline (10) | 66.5±1.97 | 0.0251* | 383.3±31.5 | 0.1275 | 69.1±2.79 | 0.0122* | 20.0±1.46 | 0.0338* |
|  | PTH (10) | 69.1±2.63 |  | 407.9±37.0 |  | 75.4±6.15 |  | 23.0±3.67 |  |
| WSB | Saline (9) | 67.9±3.57 | 0.6705 | 419.2±50.9 | 0.9384 | 73.2±5.59 | 0.8870 | 22.6±2.88 | 0.8757 |
|  | PTH (9) | 68.6±5.50 |  | 421.4±68.4 |  | 73.7±9.20 |  | 22.3±4.49 |  |

* p <0.05, bold: p<0.00625

C. **Male – 12 wks whole body BMD and Delta BMD**

|  |  | BMD (mg/cm^2^) | | Change from Baseline (%) | |
| --- | --- | --- | --- | --- | --- |
| Strain | Treatment  (N) | Average | P value | Average | P value |
| 129 | Saline | 75.70±1.61 | 0.9829 | -4.10±3.11 | 0.1048 |
|  | PTH | 75.68±1.64 |  | 0.00±6.74 |  |
| A/J | Saline | 63.12±2.04 | 0.6227 | 2.72±4.18 | 0.0140* |
|  | PTH | 63.51±1.35 |  | 8.21±4.81 |  |
| B6 | Saline (8) | 70±3.989 | 0.9194 | 7.37±6.38 | 0.1874 |
| r | PTH (10) | 70.20±4.47 |  | 11.28±5.34 |  |
| CAST | Saline (7) | 58.63±1.8 | 0.6859 | 5.68±4.66 | 0.2849 |
|  | PTH (9) | 59.08±2.59 |  | 8.22±4.32 |  |
| NOD | Saline (9) | 75.97±2.77 | 0.3055 | 2.06±5.47 | 0.1241 |
|  | PTH (10) | 77.68±2.64 |  | 5.93±4.86 |  |
| NZO | Saline (9) | 69.42±4.15 | 0.6121 | 9.73±4.22 | 0.4783 |
|  | PTH (10) | 70.33±3.75 |  | 7.56±8.71 |  |
| PWK | Saline (10) | 64.22±3.32 | 0.3799 | -2.98±6.44 | 0.3343 |
|  | PTH (10) | 65.26±1.48 |  | -0.31±5.34 |  |
| WSB | Saline (10) | 61.43±2.89 | 0.6222 | 5.39±6.0 | 0.1737 |
|  | PTH (10) | 60.57±4.59 |  | 9.43±6.71 |  |

* p <0.05, bold: p<0.00625

D. **Female – 12 wks and Delta BMD**

|  |  | BMD (mg/cm^2^) | | Change from Baseline (%) | |
| --- | --- | --- | --- | --- | --- |
| Strain | Treatment  (N) | Average | P value | Average | P value |
| 129 | Saline (10) | 74.40±4.56 | 0.8473 | -1.39±4.97 | 0.0456 |
|  | PTH (10) | 74.06±3.24 |  | 4.34±6.69 |  |
| A/J | Saline (9) | 62.19±1.92 | 0.5204 | 5.41±4.29 | 0.0598 |
|  | PTH (10) | 61.61±2.0 |  | 9.13±3.75 |  |
| B6 | Saline (10) | 60.16±2.28 | 0.1990 | 14.43±3.91 | 0.1885 |
|  | PTH (10) | 61.95±3.34 |  | 18.97±8.97 |  |
| CAST | Saline (8) | 59.0±2.06 | 0.6689 | 0.69±4.69 | 0.1062 |
|  | PTH (10) | 58.57±2.20 |  | 8.22±4.32 |  |
| NOD | Saline (10) | 70.61±4.47 | 0.0265* | 4.29±7.84 | 0.8818 |
|  | PTH (10) | 74.95±3.44 |  | 4.79±6.95 |  |
| NZO | Saline (10) | 74.99±6.93 | 0.0087* | -0.54±8.9 | **0.0058** |
|  | PTH (10) | 67.56±2.99 |  | 9.73±4.22 |  |
| PWK | Saline (10) | 65.94±3.28 | 0.5700 | 0.93±4.69 | 0.0352* |
|  | PTH (10) | 65.11±2.87 |  | 6.22±5.54 |  |
| WSB | Saline (9) | 60.93±8.51 | 0.4931 | 12.44±11.53 | 0.4978 |
|  | PTH (9) | 58.64±6.44 |  | 15.49±6.10 |  |

* p <0.05, bold: p<0.00625

**Supplemental Table 3**. Femoral Cortical Architecture

A. **Male**

|  |  | Cortical Area (mm^2^) | | Marrow Area (mm^2^) | | Cortical Thickness (mm) | | Cortical Area/  Total area | |
| --- | --- | --- | --- | --- | --- | --- | --- | --- | --- |
| Strain | Treatment | Average | P value | Average | P value | Average | P value | Average | P value |
| 129 | Saline | 0.89±0.08 | 0.0557 | 0.68±0.05 | 0.2562 | 0.232±0.018 | 0.0324* | 0.55±0.03 | 0.009* |
|  | PTH | 0.97±0.09 |  | 0.66±0.04 |  | 0.251±0.017 |  | 0.58±0.02 |  |
| A/J | Saline | 0.78±0.06 | 0.0066* | 0.52±0.04 | 0.3005 | 0.230±0.012 | **0.0050** | **0.58**±0.02 | 0.060 |
|  | PTH | 0.86±0.07 |  | 0.54±0.04 |  | 0.244±0.010 |  | 0.59±0.01 |  |
| B6 | Saline | 0.87±0.07 | 0.8608 | 1.34±0.13 | 0.0891 | 0.169±0.017 | 0.9636 | 0.38±0.02 | 0.241 |
|  | PTH | 0.96±0.09 |  | 1.47±0.18 |  | 0.169±0.013 |  | 0.37±0.03 |  |
| CAST | Saline | 0.74±0.05 | 0.0156* | 0.53±0.04 | 0.0337* | 0.217±0.011 | **0.0011** | 0.57±0.03 | **0.005** |
|  | PTH | 0.82±0.05 |  | 0.48±0.02 |  | 0.242±0.013 |  | 0.61±0.02 |  |
| NOD | Saline | 0.87±0.08 | 0.2074 | 0.70±0.05 | 0.1826 | 0.219±0.018 | 0.4933 | 0.54±0.03 | 0.829 |
|  | PTH | 0.97±0.09 |  | 0.76±0.11 |  | 0.225±0.015 |  | 0.53±0.04 |  |
| NZO | Saline | 1.06±0.08 | 0.5296 | 1.16±0.07 | 0.8290 | 0.230±0.014 | 0.9482 | 0.48±0.02 | 0.472 |
|  | PTH | 1.08±0.05 |  | 1.15±0.07 |  | 0.231±0.010 |  | 0.48±0.02 |  |
| PWK | Saline | 0.45±0.04 | **0.0011** | 0.61±0.05 | 0.4482 | 0.146±0.006 | **0.0032** | 0.43±0.01 | 0.049* |
|  | PTH | 0.52±0.04 |  | 0.63±0.06 |  | 0.163±0.013 |  | 0.45±0.03 |  |
| WSB | Saline | 0.54±0.04 | 0.6775 | 0.78±0.08 | 0.8396 | 0.155±0.011 | 0.7930 | 0.41±0.03 | 0.935 |
|  | PTH | 0.55±0.02 |  | 0.78±0.04 |  | 0.156±0.006 |  | 0.41±0.01 |  |

* p <0.05, bold: p<0.00625

B. **Female**

|  |  | Cortical Area (mm^2^) | | Marrow Area (mm^2^) | | Cortical Thickness (mm) | | Cortical Area/  Total area (%) | |
| --- | --- | --- | --- | --- | --- | --- | --- | --- | --- |
| Strain | Treatment | Average | P value | Average | P value | Average | P value | Average | P value |
| 129 | Saline | 0.84±0.07 | 0.0145* | 0.67±0.07 | 0.0168 | 0.227±0.015 | **<0.001** | 0.54±0.03 | **<0.001** |
|  | PTH | 0.92±0.06 |  | 0.60±0.04 |  | 0.250±0.011 |  | 0.59±0.02 |  |
| A/J | Saline | 0.74±0.05 | 0.1325 | 0.49±0.04 | **0.0014** | 0.224±0.010 | **<0.001** | 0.58±0.02 | **<0.001** |
|  | PTH | 0.77±0.03 |  | 0.43±0.04 |  | 0.243±0.005 |  | 0.62±0.01 |  |
| B6 | Saline | 0.75±0.06 | 0.0134* | 1.00±0.09 | 0.8680 | 0.173±0.012 | **0.0030** | 0.42±0.03 | 0.0080* |
|  | PTH | 0.83±0.08 |  | 0.96±0.06 |  | 0.192±0.016 |  | 0.45±0.03 |  |
| CAST | Saline | 0.66±0.04 | 0.1110 | 0.47±0.04 | **<0.001** | 0.206±0.010 | 0.0090* | 0.56±0.02 | **<0.001** |
|  | PTH | 0.71±0.07 |  | 0.39±0.02 |  | 0.232±0.020 |  | 0.62±0.03 |  |
| NOD | Saline | 0.78±0.03 | **0.0002** | 0.58±0.06 | 0.2862 | 0.218±0.011 | 0.0132* | 0.56±0.03 | 0.3217 |
|  | PTH | 0.85±0.04 |  | 0.60±0.04 |  | 0.231±0.010 |  | 0.57±0.02 |  |
| NZO | Saline | 0.92±0.05 | 0.1519 | 0.95±0.06 | 0.1730 | 0.222±0.011 | 0.5026 | 0.49±0.02 | 0.0343* |
|  | PTH | 0.97±0.07 |  | 0.91±0.08 |  | 0.225±0.011 |  | 0.52±0.02 |  |
| PWK | Saline | 0.53±0.04 | 0.0333* | 0.57±0.06 | 0.7849 | 0.171±0.012 | 0.0659 | 0.48±0.03 | 0.1794 |
|  | PTH | 0.57±0.04 |  | 0.58±0.03 |  | 0.180±0.007 |  | 0.50±0.01 |  |
| WSB | Saline | 0.56±0.08 | 0.3565 | 0.85±0.38 | 0.1803 | 0.157±0.025 | 0.0506 | 0.41±0.07 | 0.0434* |
|  | PTH | 0.60±0.07 |  | 0.66±0.05 |  | 0.178±0.015 |  | 0.47±0.03 |  |

* p <0.05, bold: p<0.00625

**Supplemental Table 4**. Femoral Mechanical Properties

Male

|  |  | Breaking  Strength | | Stiffness (N/m) | | Work to Failure (kJ) | |
| --- | --- | --- | --- | --- | --- | --- | --- |
| Strain | Treatment | Average | P value | Average | P value | Average | P value |
| 129 | Saline | 24.33±2.93 | 0.2517 | 119.43±12.82 | 0.2158 | 5.57±0.94 | 0.4164 |
|  | PTH | 25.90±2.81 |  | 127.32±13.94 |  | 6.24±2.35 |  |
| A/J | Saline | 16.82±2.6 | **0.0036** | 88.86±17.44 | 0.3043 | 5.39±1.57 | 0.6369 |
|  | PTH | 20.31±1.97 |  | 95.68±10.32 |  | 5.06±1.50 |  |
| B6 | Saline | 18.57±1.83 | 0.0726 | 89.47±16.19 | 0.4680 | 15.96±4.68 | 0.7704 |
|  | PTH | 20.30±2.21 |  | 93.72±7.76 |  | 15.35±4.54 |  |
| CAST | Saline | 18.57±2.50 | 0.0653 | 80.79±18.30 | 0.1028 | 8.60±2.37 | 0.7202 |
|  | PTH | 20.84±1.70 |  | 95.41±13.11 |  | 8.95±0.88 |  |
| NOD | Saline | 20.18±1.46 | 0.0532 | 95.87±10.72 | 0.0343* | 6.50±1.46 | 0.1420 |
|  | PTH | 21.96±2.26 |  | 107.03±11.08 |  | 7.75±2.08 |  |
| NZO | Saline | 31.15±2.63 | 0.5139 | 142±20.24 | 0.4328 | 8.83±2.77 | 0.5700 |
|  | PTH | 31.78±0.98 |  | 148.72±7.03 |  | 8.19±1.90 |  |
| PWK | Saline | 13.04±0.96 | **0.0042** | 66.55±8.89 | 0.1159 | 4.85±1.77 | 0.6178 |
|  | PTH | 15.75±2.24 |  | 73.17±8.43 |  | 5.22±1.39 |  |
| WSB | Saline | 15.87±1.24 | 0.2439 | 75.73±8.48 | 0.0518 | 9.42±2.07 | 0.6039 |
|  | PTH | 16.60±1.37 |  | 85.10±10.97 |  | 9.90±1.29 |  |

* p <0.05, bold: p<0.00625

Female

|  |  | Breaking  Strength | | Stiffness (N/m) | | Work to Failure (kJ) | |
| --- | --- | --- | --- | --- | --- | --- | --- |
| Strain | Treatment |  |  | Average | P value | Average | P value |
| 129 | Saline | 22.59±2.92 | 0.1634 | 122.64±12.97 | 0.7607 | 5.24±0.83 | 0.3307 |
|  | PTH | 24.20±1.87 |  | 124.81±17.95 |  | 6.07±2.43 |  |
| A/J | Saline | 16.93±1.68 | 0.6271 | 88.02±12.79 | 0.7925 | 4.76±2.33 | 0.7387 |
|  | PTH | 17.29±1.55 |  | 89.44±10.89 |  | 4.45±1.82 |  |
| B6 | Saline | 15.23±1.35 | **0.0025** | 76.18±10.16 | 0.0395* | 11.43±2.87 | 0.5554 |
|  | PTH | 18.16±2.18 |  | 87.57±12.59 |  | 12.41±4.26 |  |
| CAST | Saline | 16.64±2.84 | 0.7618 | 70.11±20.41 | 0.5056 | 7.20±3.02 | 0.6038 |
|  | PTH | 17.04±2.67 |  | 76.90±21.44 |  | 6.62±1.50 |  |
| NOD | Saline | 17.60±1.44 | 0.0292* | 84.83±12.56 | 0.0740 | 6.24±0.91 | 0.6600 |
|  | PTH | 19.30±1.67 |  | 96.14±10.71 |  | 6.48±1.40 |  |
| NZO | Saline | 26.10±2.19 | 0.9266 | 135.12±10.73 | 0.1706 | 7.67±2.34 | 0.4588 |
|  | PTH | 26.01±2.21 |  | 124.05±21.65 |  | 6.86±2.46 |  |
| PWK | Saline | 14.24±1.73 | 0.5253 | 77.84±10.05 | 0.9834 | 3.83±1.27 | 0.7193 |
|  | PTH | 14.70±1.43 |  | 77.73±12.94 |  | 3.61±1.42 |  |
| WSB | Saline | 15.68±2.38 | 0.3742 | 83.46±16.86 | 0.7153 | 7.40±2.47 | 0.2636 |
|  | PTH | 16.74±2.54 |  | 76.85±50.11 |  | 8.68±2.21 |  |

* p <0.05, bold: p<0.00625

**Supplemental Table 5**. Femoral Trabecular Architecture

A. Male

|  |  | BV/TV | | Trabecular Thickness | | Trabecular Number | |
| --- | --- | --- | --- | --- | --- | --- | --- |
| Strain | Treatment | Average | P value | Average | P value | Average | P value |
| 129 | Saline | 0.066±0.020 | 0.4233 | 0.044±0.006 | 0.2742 | 4.00±0.21 | 0.6445 |
|  | PTH | 0.075±0.026 |  | 0.046±0.006 |  | 3.94±0.34 |  |
| A/J | Saline | 0.040±0.013 | 0.0189* | 0.046±0.002 | 0.0088* | 2.91±0.27 | 0.0124* |
|  | PTH | 0.055±0.012 |  | 0.050±0.003 |  | 3.21±0.19 |  |
| B6 | Saline | 0.126±0.021 | 0.3302 | 0.050±0.003 | **0.0003** | 4.43±0.38 | **0.0024** |
|  | PTH | 0.140±0.036 |  | 0.061±0.006 |  | 3.49±0.53 |  |
| CAST* | Saline | 0.017±0.006 | **0.0003** | 0.048±0.006 | 0.0903 |  |  |
|  | PTH | 0.004±0.003 |  | 0.037±0.014 |  |  |  |
| NOD | Saline | 0.059±0.013 | **0.0014** | 0.060±0.004 | 0.8131 | 2.55±0.31 | 0.0100* |
|  | PTH | 0.029±0.008 |  | 0.060±0.007 |  | 2.06±0.15 |  |
| NZO | Saline | 0.093±0.022 | 0.5899 | 0.051±0.005 | 0.0079* | 3.68±0.36 | 0.0288* |
|  | PTH | 0.088±0.019 |  | 0.064±0.011 |  | 3.28±0.37 |  |
| PWK | Saline | 0.044±0.010 | 0.5970 | 0.040±0.022 | 0.0151* | 2.64±0.35 | 0.8034 |
|  | PTH | 0.047±0.015 |  | 0.043±0.003 |  | 2.61±0.27 |  |
| WSB | Saline | 0.053±0.015 | 0.1878 | 0.048±0.002 | 0.3954 | 2.96±0.32 | 0.3508 |
|  | PTH | 0.046±0.008 |  | 0.049±0.005 |  | 2.82±0.32 |  |

*Trabecular number could not be calculated for the CAST mice as the value was below the lower limit of detection of the µCT instrument, * p <0.05, bold: p<0.00625

B. Female

|  |  | BV/TV | | Trabecular Thickness | | Trabecular Number | |
| --- | --- | --- | --- | --- | --- | --- | --- |
| Strain | Treatment | Average | P value | Average | P value | Average | P value |
| 129 | Saline | 0.093±0.026 | **<0.001** | 0.043±0.003 | **<0.001** | 4.62±0.41 | 0.1309 |
|  | PTH | 0.171±0.030 |  | 0.051±0.003 |  | 4.88±0.30 |  |
| A/J | Saline | 0.054±0.022 | 0.3099 | 0.047±0.004 | 0.1531 | 3.04±0.34 | 0.4562 |
|  | PTH | 0.064±0.018 |  | 0.049±0.002 |  | 3.15±0.28 |  |
| B6 | Saline | 0.041±0.009 | 0.1858 | 0.048±0.004 | 0.5722 | 3.02±0.27 | 0.1950 |
|  | PTH | 0.047±0.010 |  | 0.049±0.005 |  | 3.22±0.29 |  |
| CAST* | Saline | 0.015±0.007 | 0.0209* | 0.048±0.007 | 0.7799 |  |  |
|  | PTH | 0.007±0.005 |  | 0.047±0.015 |  |  |  |
| NOD | Saline | 0.054±0.018 | 0.5473 | 0.056±0.006 | 0.0328* | 2.47±0.42 | 0.6837 |
|  | PTH | 0.059±0.015 |  | 0.063±0.006 |  | 2.54±0.30 |  |
| NZO | Saline | 0.175±0.033 | 0.2531 | 0.058±0.002 | **0.0012** | 4.20±0.33 | 0.2591 |
|  | PTH | 0.190±0.023 |  | 0.065±0.005 |  | 4.06±0.17 |  |
| PWK | Saline | 0.153±0.033 | 0.1075 | 0.046±0.003 | 0.0357* | 5.01±0.38 | 0.9640 |
|  | PTH | 0.175±0.018 |  | 0.049±0.002 |  | 5.00±0.38 |  |
| WSB | Saline | 0.041±0.012 | 0.0478* | 0.046±0.004 | 0.9637 | 2.72±0.34 | 0.0281* |
|  | PTH | 0.060±0.021 |  | 0.046±0.005 |  | 3.54±0.90 |  |

*Trabecular number could not be calculated for the CAST mice as the value was below the lower limit of detection of the µCT instrument, * p <0.05, bold: p<0.00625

**Supplemental Table 6**. Vertebral Trabecular Architecture

A. Male

|  |  | BV/TV | | Trabecular Thickness | | Trabecular Number | |
| --- | --- | --- | --- | --- | --- | --- | --- |
| Strain | Treatment | Average | P value | Average | P value | Average | P value |
| 129 | Saline | 0.170±0.021 | 0.2646 | 48.2±3.11 | 0.1448 | 4.21±0.25 | 0.8866 |
|  | PTH | 0.186±0.037 |  | 50.8±3.90 |  | 4.23±0.41 |  |
| A/J | Saline | 0.134±0.012 | **<0.001** | 48.3±2.32 | 0.0267* | 3.52±0.23 | 0.1132 |
|  | PTH | 0.173±0.019 |  | 52.2±4.09 |  | 3.79±0.41 |  |
| B6 | Saline | 0.192±0.037 | 0.4530 | 49.4±4.40 | 0.0293* | 5.19±0.29 | **0.0044** |
|  | PTH | 0.203±0.034 |  | 53.4±3.64 |  | 4.69±0.51 |  |
| CAST | Saline | 0.072±0.007 | 0.4272 | 40.1±2.34 | 0.9547 | 3.27±0.09 | **0.0008** |
|  | PTH | 0.069±0.011 |  | 40.2±3.15 |  | 2.90±0.22 |  |
| NOD | Saline | 0.140±0.020 | 0.0233* | 53.1±1.62 | **<0.001** | 3.49±0.38 | 0.1254 |
|  | PTH | 0.161±0.015 |  | 62.3±3.20 |  | 3.25±0.26 |  |
| NZO | Saline | 0.179±0.018 | **<0.001** | 50.2±2.05 | **<0.001** | 4.28±0.23 | 0.1600 |
|  | PTH | 0.237±0.021 |  | 60.5±3.50 |  | 4.13±0.21 |  |
| PWK | Saline | 0.154±0.013 | **<0.001** | 42.2±1.32 | **<0.001** | 4.26±0.24 | 0.4497 |
|  | PTH | 0.201±0.014 |  | 47.3±1.16 |  | 4.34±0.22 |  |
| WSB | Saline | 0.188±0.032 | 0.2184 | 48.6±2.17 | 0.2499 | 4.53±0.32 | 0.6616 |
|  | PTH | 0.207±0.031 |  | 49.9±2.69 |  | 4.61±0.40 |  |

* p <0.05, bold: p<0.00625

B. Female

|  |  | BV/TV | | Trabecular Thickness | | Trabecular Number | |
| --- | --- | --- | --- | --- | --- | --- | --- |
| Strain | Treatment | Average | P value | Average | P value | Average | P value |
| 129 | Saline | 0.281±0.039 | 0.2502 | 55.2±4.78 | 0.0270* | 5.02±0.52 | 0.5639 |
|  | PTH | 0.261±0.037 |  | 50.9±2.77 |  | 5.13±0.33 |  |
| A/J | Saline | 0.159±0.029 | 0.5760 | 52.6±1.35 | 0.3684 | 3.16±0.33 | 0.6274 |
|  | PTH | 0.153±0.019 |  | 48.5±13.64 |  | 3.29±0.72 |  |
| B6 | Saline | 0.160±0.043 | 0.9243 | 48.3±4.80 | 0.3742 | 4.22±0.64 | 0.1474 |
|  | PTH | 0.162±0.025 |  | 50.0±2.34 |  | 3.88±0.19 |  |
| CAST | Saline | 0.080±0.014 | 0.1207 | 38.6±1.71 | 0.0196* | 3.50±0.52 | 0.5630 |
|  | PTH | 0.096±0.24 |  | 42.5±3.63 |  | 3.42±0.25 |  |
| NOD | Saline | 0.133±0.026 | **0.0043** | 52.6±4.22 | **<0.001** | 2.91±0.33 | 0.3391 |
|  | PTH | 0.176±0.029 |  | 60.7±2.60 |  | 3.06±0.35 |  |
| NZO | Saline | 0.282±0.020 | 0.3548 | 56.6±3.95 | 0.9592 | 4.86±0.19 | 0.3940 |
|  | PTH | 0.264±0.055 |  | 56.5±4.65 |  | 4.72±0.46 |  |
| PWK | Saline | 0.226±0.020 | 0.0067* | 48.4±1.71 | 0.0101* | 4.52±0.22 | 0.8456 |
|  | PTH | 0.259±0.026 |  | 51.0±2.61 |  | 4.55±0.42 |  |
| WSB | Saline | 0.191±0.033 | 0.0300* | 49.4±4.88 | 0.1158 | 4.51±0.11 | 0.1729 |
|  | PTH | 0.233±0.041 |  | 53.0±4.15 |  | 4.38±0.24 |  |

* p <0.05, bold: p<0.00625
